# Coevolutionary phage training leads to greater bacterial suppression and delays the evolution of phage resistance

**DOI:** 10.1101/2020.11.02.365361

**Authors:** Joshua M. Borin, Sarit Avrani, Jeffrey E. Barrick, Katherine L. Petrie, Justin R. Meyer

## Abstract

The evolution of antibiotic resistant bacteria threatens to become the leading cause of worldwide mortality. This crisis has renewed interest in the practice of phage therapy. Yet, bacteria’s capacity to evolve resistance is likely to debilitate this therapy as well. To combat the evolution of phage resistance and improve treatment outcomes, many have suggested leveraging phages’ ability to counter resistance by evolving phages on target hosts before using them in therapy (phage training). We found that during in vitro experiments, a phage trained for 28 days suppressed bacteria ∼1000-fold for 3-8 times longer than its untrained ancestor. This extension was due to a delay in the evolution of resistance. Several factors contributed to this prolonged suppression. Mutations that confer resistance to trained phages are ∼100× less common and, while the target bacterium can evolve complete resistance to the untrained phage in a single step, multiple mutations are required to evolve complete resistance to trained phages. Mutations that confer resistance to trained phages are more costly than mutations for untrained phage resistance. And when resistance does evolve, trained phages are better able to suppress these forms of resistance. One way the trained phage improved was through recombination with a gene in a defunct prophage in the host genome, which doubled phage fitness. This direct transfer of information encoded by the host but originating from a relict phage provides a previously unconsidered mode of training phage. Overall, we provide a case study for successful phage training and uncover mechanisms underlying its efficacy.

**Significance Statement:** The evolution of antibiotic resistant bacteria threatens to claim over 10 million lives annually by 2050. This crisis has renewed interest in phage therapy, the use of bacterial viruses to treat infections. A major barrier to successful phage therapy is that bacteria readily evolve phage resistance. One idea proposed to combat resistance is “training” phages by using their natural capacity to evolve to counter resistance. Here, we show that training phages by coevolving them with their host for one month enhanced their capacity for suppressing bacterial growth and delayed the emergence of resistance. Enhanced suppression was caused by several mechanisms, suggesting that the coevolutionary training protocol produces a robust therapeutic that employs complementary modes of action.

## Introduction

In 30 years, the World Health Organization predicts that antibiotic resistant bacteria will kill over 10 million people each year—more deaths than that caused by cancer (1). This health crisis, in part caused by the heavy and often inappropriate way we use antibiotic drugs, has led to the spread of resistance genes through clinical and natural environments and to the emergence of multidrug resistant (MDR) “superbugs” that are often untreatable due to their resistance against all available classes of antibiotics (2-4). As bacteria continue to outpace our discovery and development of new drugs, the evolution of resistance threatens to return us to a pre-antibiotic era of infectious disease (5,6).

This crisis has renewed interest in the century-old practice of phage therapy: the use of phages, viruses that infect bacteria, to treat bacterial infections (7-11). Recently, phage therapy has shown promise in cases where drugs of last resort fail to treat life-threatening MDR bacterial infections (9-13). However, even in successful cases, the evolution of *phage* resistance poses a considerable threat to the efficacy of treatment (9,12,14). For example, in 2016, at the University of California San Diego, a patient with acute pancreatitis complicated by an MDR *Acinetobacter baumanii* infection was treated with two 4-phage cocktails that suppressed the pathogen in vitro (9). Within 8 days, *A. baumanii* isolated from the patient was resistant to all 8 phages used. Fortunately, the infection resolved following delivery of a ninth phage and the patient survived. This case is representative of numerous phage therapy studies (9-12,14). A metanalysis in 2018 reported that phage resistance evolved in 82% of animal gut decolonization studies, 50% of meningitis/sepsis models, and 75% of human clinical cases where the evolution of resistance was monitored (14). These observations of rapid phage resistance evolution in therapy mirror decades of basic research in the lab; mutations that confer resistance to phages are often as common as those for antibiotic resistance (15-17). Furthermore, many of these resistance mutations confer cross-resistance to multiple phages (18).

Although resistance to phages is as or more common than to antibiotics, advantages of using phages as therapeutics have been proposed time and again (7,8,19-22). Notably, unlike antibiotics, phages are biological entities that evolve. By reciprocally adapting to changes in their hosts (coevolution), phages have maintained the ability to infect their hosts for millennia. Many have proposed harnessing this inherent, evolutionary potential by preemptively coevolving phages with target bacterial prey (22-24). Proponents of this “phage training” approach suggest that, by experiencing the ways their host can evolve resistance, trained phages will evolve to counter host defenses. Then, trained phages “from the future” can be used to trap the ancestral, un-coevolved bacteria from their past that are infecting the patient.

While the idea of phage training is enticing, it has not yet been adopted for therapy. Contrasting theories of bacteria-phage coevolutionary dynamics make the success of phage training uncertain (12,24,25). According to some conceptual models of coevolution (e.g. matching alleles), as phages adapt to their evolving host, they lose the ability to infect past hosts (24). In such cases, phage training would not work because trained phages would lose the ability to infect the original target bacterium. Alternatively, other models of coevolution (e.g. gene for gene, arms race dynamics) argue that as phages adapt to their evolving host, they maintain the ability to infect their original host (24). In this scenario, training would expand phage host range to encompass both original and contemporary bacteria. Regardless of how coevolution affects host range, some opponents of phage training contend that the use of trained phages will apply stronger selection on target bacteria which will accelerate the evolution of resistance and loss of therapeutic efficacy (22).

In this study, we conducted a coevolution experiment using *Escherichia coli* and either untrained or trained phages to evaluate the potential of phage training for therapy. By comparing the population dynamics of coevolving bacteria and phages, we find that trained phages suppress the target bacteria more strongly and for longer than untrained phages. Through post hoc analyses on the bacteria and phages that evolved in our experiment, we identify the factors that allowed trained phages to suppress host populations and delay the evolution of resistance.

## Results

### Coevolution Experiment

We propagated populations of *E. coli* B strain REL606 with either untrained or trained strains of phage λ in flasks for 30 days and compared the population dynamics of coevolving bacteria and phages. For untrained phages, we used λunt, a lytic strain of λ that uses LamB as a receptor. For trained phages, we used λtrn, a descendant of λunt that was isolated on day 28 of a previous coevolution experiment with REL606 and that can infect using either of two receptors, LamB and OmpF (26,35). Because λtrn uses two receptors, we hypothesized that it would be more difficult for hosts to evolve envelope-based resistance.

Daily measurements of population densities revealed large differences in bacterial abundance between treatments (Fig. 1A), whereas phage densities were similar (Fig. 1B). By day 3, bacteria coevolving with λunt were no longer suppressed by phages and were instead limited by the availability of glucose in the media. This rapid loss of suppression is consistent with past studies on λunt and REL606 (26-28), as well as studies on other bacteria-phage pairs (29,30); when cultured with phages in flasks, bacteria rapidly evolve resistance and their growth is limited by the availability of nutrients. However, in flasks with λtrn, phages suppressed their hosts for 13-28 days, at which point bacterial densities slowly increased until they became limited by the availability of glucose, as in flasks with λunt. The prolonged suppression of bacteria by λtrn suggested that resistance to λtrn evolved much later than to λunt.

**Figure 1.**
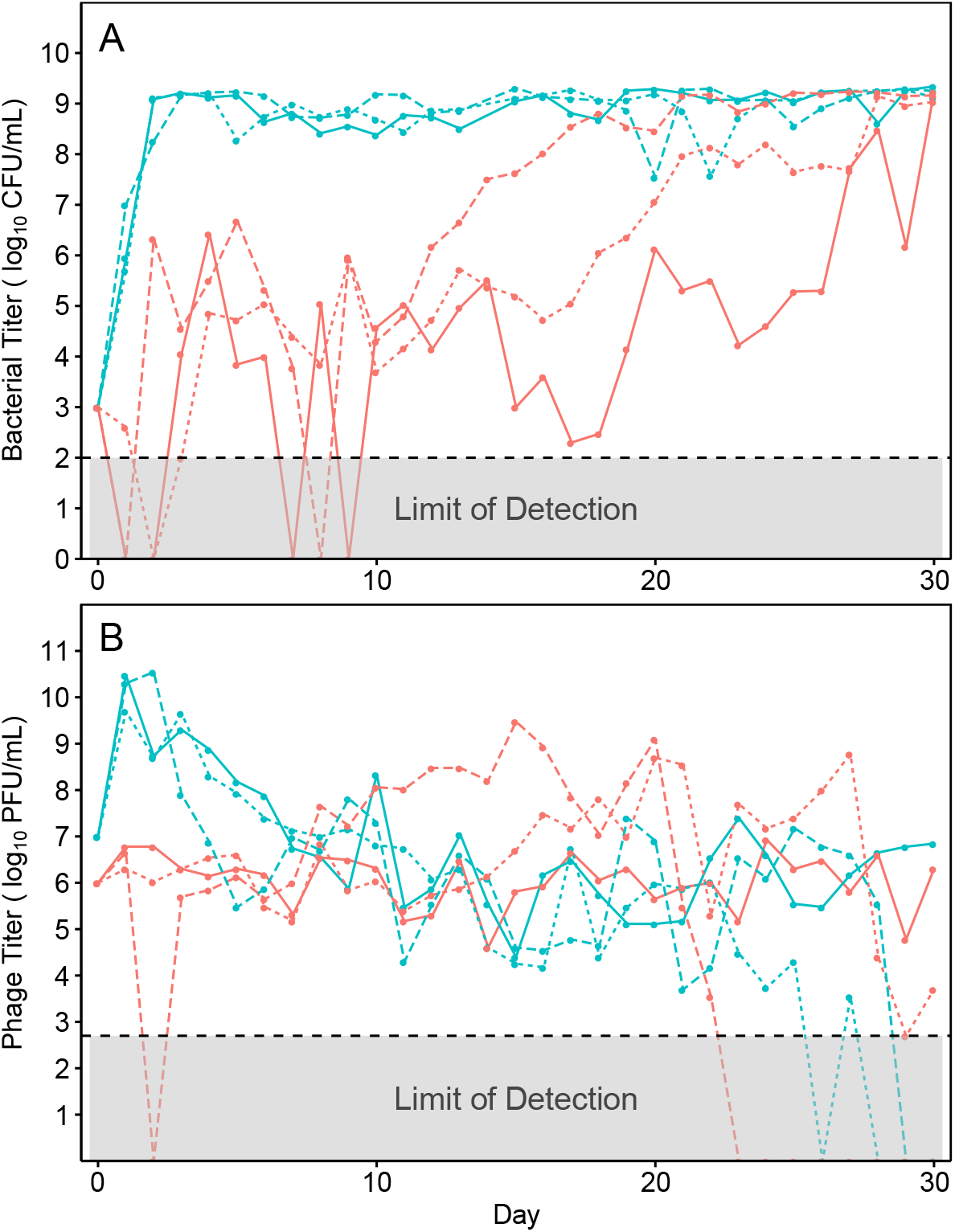
Population dynamics of bacteria (A) and phage (B) during 30 days of coevolution in flasks, estimated from colony forming units (CFU) and plaque forming units (PFU), respectively. Flasks containing λunt are teal and λtrn are red. Line types correspond with replicate Populations (solid=1, dotted=2, dashed=3).

### Emergence of Resistance

To determine when resistance evolved in each treatment, we sampled bacteria from various timepoints and challenged them with the initial phage they were co-cultured with (λunt or λtrn). We assessed the resistance of 12 isolates every 5 days to quantify the diversity of resistance. Initially, we performed spot assays where 5 μL of phage lysate was spotted on top of an isogenic lawn of bacteria growing on an agar plate. We found that this approach was inadequate because many bacterial isolates appeared sensitive to λtrn when grown on agar (clearing due to phage killing) but were able to grow with phage in liquid culture.

Given that the experiments were conducted in liquid medium, we developed a protocol to efficiently track bacterial population growth in liquid culture using a microtiter plate reader. Resistance was categorized visually by comparing the difference in cell density (via optical density; OD) dynamics between cultures growing with and without phage. Bacteria were deemed sensitive if no growth was observed in the presence of phage and completely resistant if cells grew uninhibited by phage. Partial resistance was recorded for bacteria that showed signs of growth but were clearly inhibited by the phage (representative isolates in Fig. S1).

Using the liquid-based assay described above, we characterized the proportion of resistance in populations of the coevolution experiment at various time-points (Fig. 2, A-F). Resistance to λunt evolved much earlier than to λtrn; by day 3, ≥50% of isolates were partially or completely resistant to λunt and by day 10, 100% of isolates were completely resistant. Despite high levels of resistance, λunt was able to persist. This was likely due to genetically “leaky resistance”, in which a resistant population of bacteria continuously generates a small number of sensitive mutant cells upon which a phage population can subsist (31). As in previous coevolution experiments, λunt in populations 1, 2, and 3 evolved to use OmpF as a receptor on days 17, 22, and 19, respectively.

**Figure 2.**
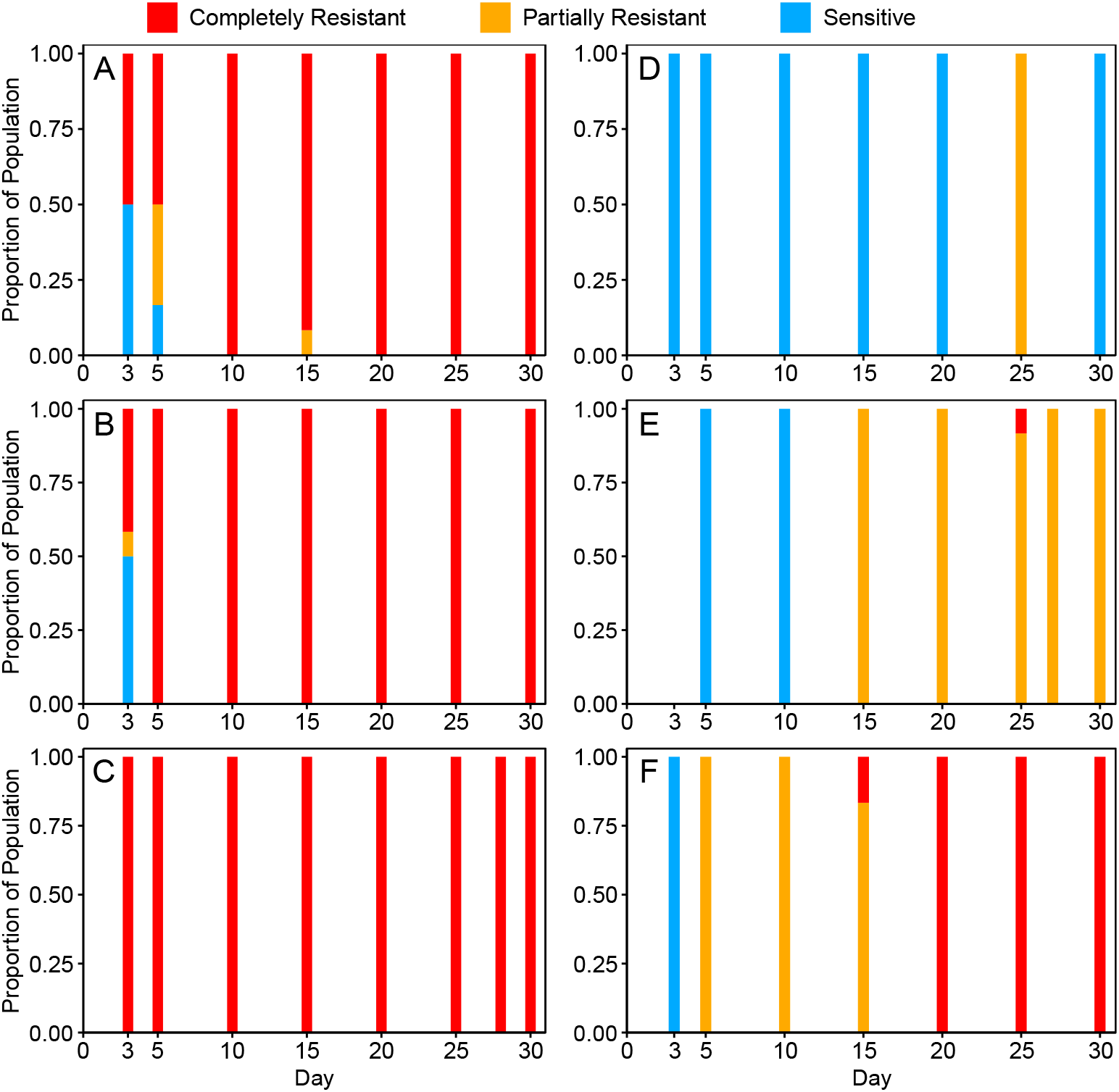
Proportion of sensitive (blue), partial (yellow) and complete (red) resistance of bacterial populations at various days of the coevolution experiment to ancestral phages from respective treatments (n=12 isolates per time-point). λunt Populations 1-3 correspond to Panels A-C and λtrn Populations 1-3 correspond to Panels D-F, respectively.

In contrast to flasks with λunt, the evolution of partial and complete resistance to λtrn occurred later and was more variable. In λtrn Population 3, partial resistance evolved by day 5 but complete resistance was not detected until day 15 (Fig. 2F). However, phages isolated 5 days later were able to infect these completely resistant isolates (Fig. S2). In λtrn Population 2, partial resistance was not detected until day 15. One completely resistant strain was isolated from day 25 but it was not detected at later time-points. Phages isolated two days later were able to infect this completely resistant isolate (Fig. S2). These results suggest that °trn’s ability to evolve counter-defenses may have contributed to the disappearance of the resistant genotype. Lastly, in λtrn Population 1, partial resistance was not detected until day 25 and complete resistance never evolved (Fig. 2D). Surprisingly, even at the end of the experiment, day-30 isolates from λtrn Population 1 lacked resistance, as determined by spot and liquid-based assays (Fig. 2D). We further describe this case in *SI Appendix:* λtrn Resistance.

Delay in the evolution of resistance against our trained phage explains the difference in suppression between flasks containing λunt and λtrn. To understand the reasons for this delay, we investigated four non-mutually exclusive hypotheses. First, λtrn resistance would take longer to evolve if there were fewer mutations available that confer resistance to this phage. Second, resistance would be delayed if more mutations were required to confer resistance to λtrn than λunt. Third, if mutations for °trn resistance are more costly, it will take them longer to rise in frequency. And fourth, λtrn may be better able to evolve counter-defenses, which would prolong its ability to suppress bacterial resistance.

### Mutation Rates of Resistance

In order to determine whether there are fewer accessible mutations for λtrn resistance than for λunt resistance, we conducted Luria-Delbruck fluctuation tests. This protocol is used to estimate the rate of mutations that generate a particular phenotype, like resistance, by applying a selective screen and enumerating cells with the desired phenotype (17,32). Because traditional fluctuation tests on agar plates could not detect some forms of resistance that evolved in our experiments (see Emergence of Resistance), we used a fluctuation test design suitable for liquid culture. Using the P0 method (32), we estimated a mutation rate for λunt resistance of 6.3 × 10^−6^ per cell. With λtrn, we found that the mutation rate for resistance was 2.2 × 10^−8^ per cell, ∼100× lower than resistance to λunt. Notably, resistance to λunt appeared to confer complete resistance, whereas λtrn resistance was only partial. Thus, mutations that confer λtrn resistance are ∼100× less common than mutations for λunt resistance, and they only confer partial resistance.

Our inability to detect mutants with complete λtrn resistance in fluctuation tests led us to focus on the second hypothesis, that complete resistance might require multiple mutations. To test this, we repeated fluctuation tests on 4 genetically distinct, partially resistant mutants isolated from the initial fluctuation test. We found that, across 10 replicate cultures for each isolate (40 total), only one replicate evolved complete resistance (with an additional mutation). Thus, under the artificial selection in fluctuation tests, multiple mutations were required to achieve complete λtrn resistance. This led us to wonder whether complete resistance to λtrn required multiple mutations in the coevolution experiment as well.

### Genetics of Resistance

To identify the resistance mutations that evolved in the coevolution experiment, we sequenced whole genomes of representative isolates at the earliest time-point that resistance was detected, as well as of isolates at later time-points with distinct changes in resistance status (i.e. greater resistance or sensitivity; Fig. S1). We identified mutations using the computational pipeline *breseq* (33) and then classified putative resistance mutations as those that occurred in genes 1) known to be involved in λ infection (26,34) or 2) that evolved in parallel in multiple populations, fluctuation tests, or previous coevolution experiments (26,35). In total, we found 2 unique putative resistance mutations across 4 bacterial isolates from flasks with λunt and 8 unique putative resistance mutations across 8 isolates from flasks with λtrn (Fig. S3).

From flasks with λunt, all completely resistant day-3 isolates had 25 bp frameshift duplications in *malT*, a positive regulator of the receptor LamB, and no other putative resistance mutations (Fig. S3). Notably, isolates with partial λunt resistance only had point mutations in *malT* (e.g. isolate P1-T5-5-P, Fig. S1C, Fig. S3). Thus, whereas frame-shift mutations in *malT* conferred complete resistance to λunt, point mutations in *malT* were less protective. *malT* appeared to be the only locus to evolve resistance mutations in the °unt treatment.

In contrast, putative resistance mutations identified in the λtrn treatment occurred at numerous loci. Many of the genes were not previously known to affect λ infection; however, they were flagged because they evolved in parallel. For example, we found a 21,535 bp 27-gene deletion in all isolates from Population 2, as well as in a fluctuation test isolate (JB43). Moreover, JB42, an independently evolved partially resistant fluctuation test isolate, had a single mutation in *lpcA*, which occurs within this 21,535 bp range. Thus, we believe the large deletion observed in Population 2 isolates confers partial resistance by removing *lpcA*. The function of the gene suggests a mechanism for partial resistance; *lpcA* codes for a phosphoheptose isomerase and is involved in assembly of the lipopolysaccharide (LPS) core and outer membrane biogenesis. It is possible that the disruption of *lpcA* alters the organization of outer membrane proteins like LamB and OmpF, which may interfere with phage adsorption. This hypothesis is also supported by the detection of parallel evolution in *waa* genes which are also associated with LPS core biogenesis. In both isolates from Population 3, we found a 4,894 bp IS1-mediated deletion of genes *waaC* through *waaT* and a nearby locus, *waaO*, evolved mutations in a previous experiment (36,37). Previously, mutations in *waa* have only been associated with phage resistance insofar as those phages use LPS as a receptor (36,37), however here they also appear to interfere with phages that use outer membrane proteins like LamB and OmpF. The mechanisms by which other putative resistance mutations confer partial λtrn resistance are less clear and are discussed in *SI Appendix:* λtrn Resistance.

After identifying putative resistance mutations that occurred in each bacterial isolate, we sought to quantify how these mutations affected phage resistance. Because strains possessed various compositions of mutations, we were unable to analyze their effects in isolation. Therefore, we conducted linear regressions to determine whether there was a relationship between the number of putative resistance mutations and the level of resistance to λunt or λtrn. To discriminate between the resistance of different isolates (especially partially resistant isolates) we needed a way to quantitatively score resistance. We tried simple approaches to score resistance by analyzing the OD trajectories of cultures (e.g. computing the difference in the area under growth curves for bacteria without and with phage), however these methods were unable to characterize differences that were easily identifiable with the naked eye. When these approaches did not work, we used principal component analysis (PCA) on the growth trajectory data, which was able to transform and score growth trajectories along a continuous scale for resistance. As a result, principal component 1 (PC1, 92.25% of variance) was able to discriminate sensitive, partial, and complete resistance, as well as between different levels of partial resistance (Fig. S4).

Using PC1 as an indicator of resistance, we conducted linear regressions to test the relationship between the number of putative resistance mutations and phenotypic resistance to ancestral phages (Fig. 3). We find that the relationship is significant for both λunt (p=0.002) and λtrn (p=0.003). The significant relationship for λtrn suggests that the putative resistance mutations we identified played a role in building bacterial defenses. This observation combined with the fluctuation test analysis provides evidence in support of the second hypothesis that multiple mutations are required to confer complete resistance to λtrn. All strains with complete resistance to λtrn possessed five putative resistance mutations (Fig. 3B), whereas isolates with complete resistance to λunt had only one mutation (Fig. 3A).

**Figure 3.**
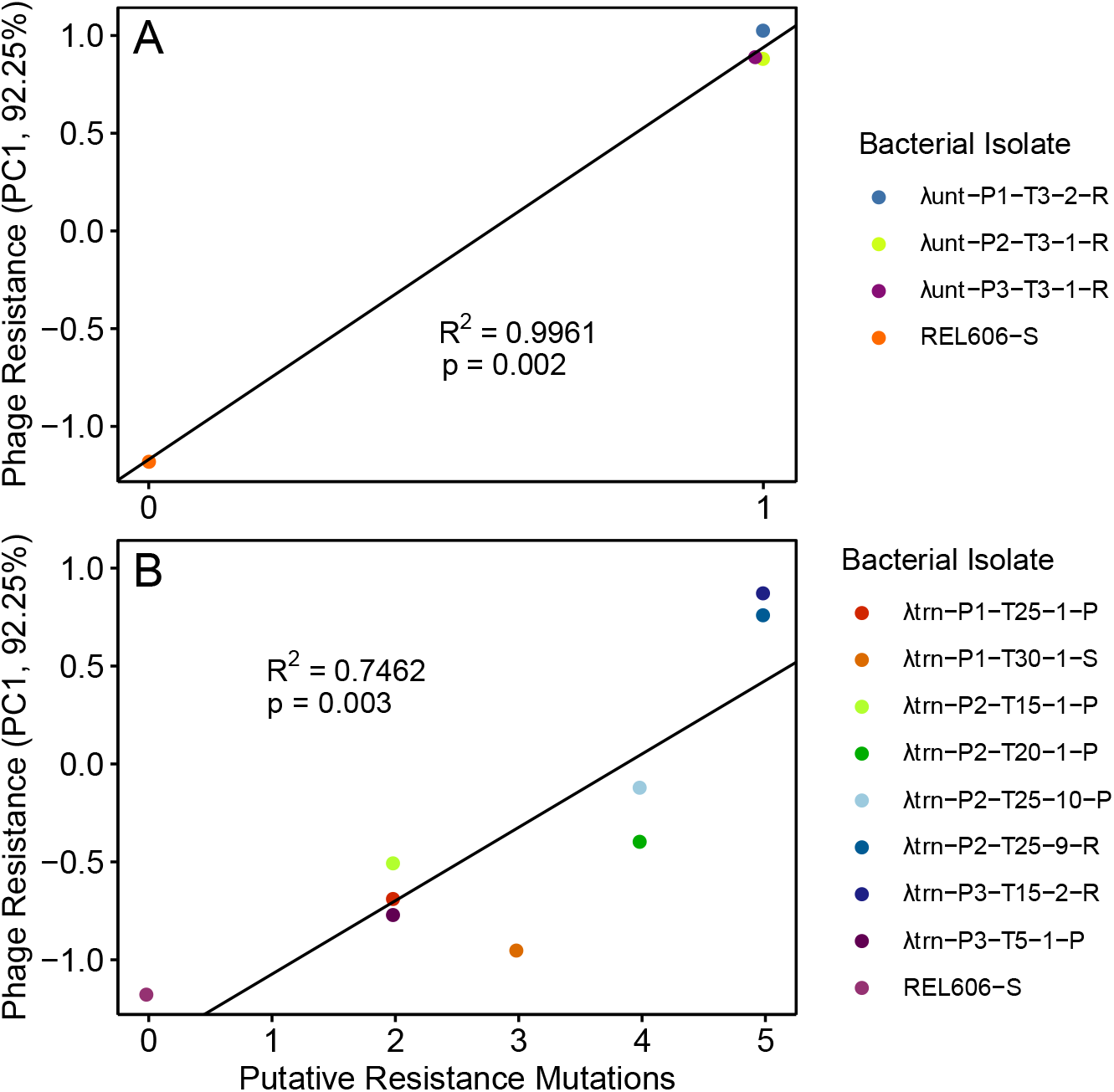
Relationship between the number of putative resistance mutations and the level of phage resistance (PC1 is indicative of resistance, see Fig. S4) from coevolutionary treatments with λunt (A) and λtrn (B). Linear regressions (PC1 ∼ Putative Resistance Mutations, black lines) are significant for both Panels A and B, however the number of mutations required for high levels of resistance is greater in B. Bacterial strains are labeled as [Treatment]-[Population]-[Day Isolated]-[Isolate #]-[Resistance Status] where S = Sensitive, R = Complete Resistance, and P = Partial Resistance to phages from respective treatments.

### Costs of Resistance

In order to determine whether resistance to λtrn was costlier than to λunt, we conducted competition experiments where resistant bacteria with different genotypes were paired with a genetically marked sensitive strain (Fig. 4). λunt resistance carried only a small cost compared to the marked competitor (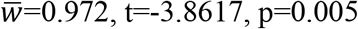, one-sample t-test). This was expected because *malT* mutants cultured in similar laboratory conditions were not found to bear a cost in previous studies (38,39). In contrast, resistance to λtrn was more costly than λunt resistance (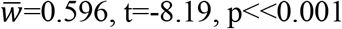, two-sample t-test). The fitness burden was similar regardless of whether the mutations conferred partial or complete resistance (p = 0.67, two-sample t-test). Thus, any level of resistance to λtrn has a cost, which helps explain the lag in the development of resistance to this phage (Fig. 1A).

**Figure 4.**
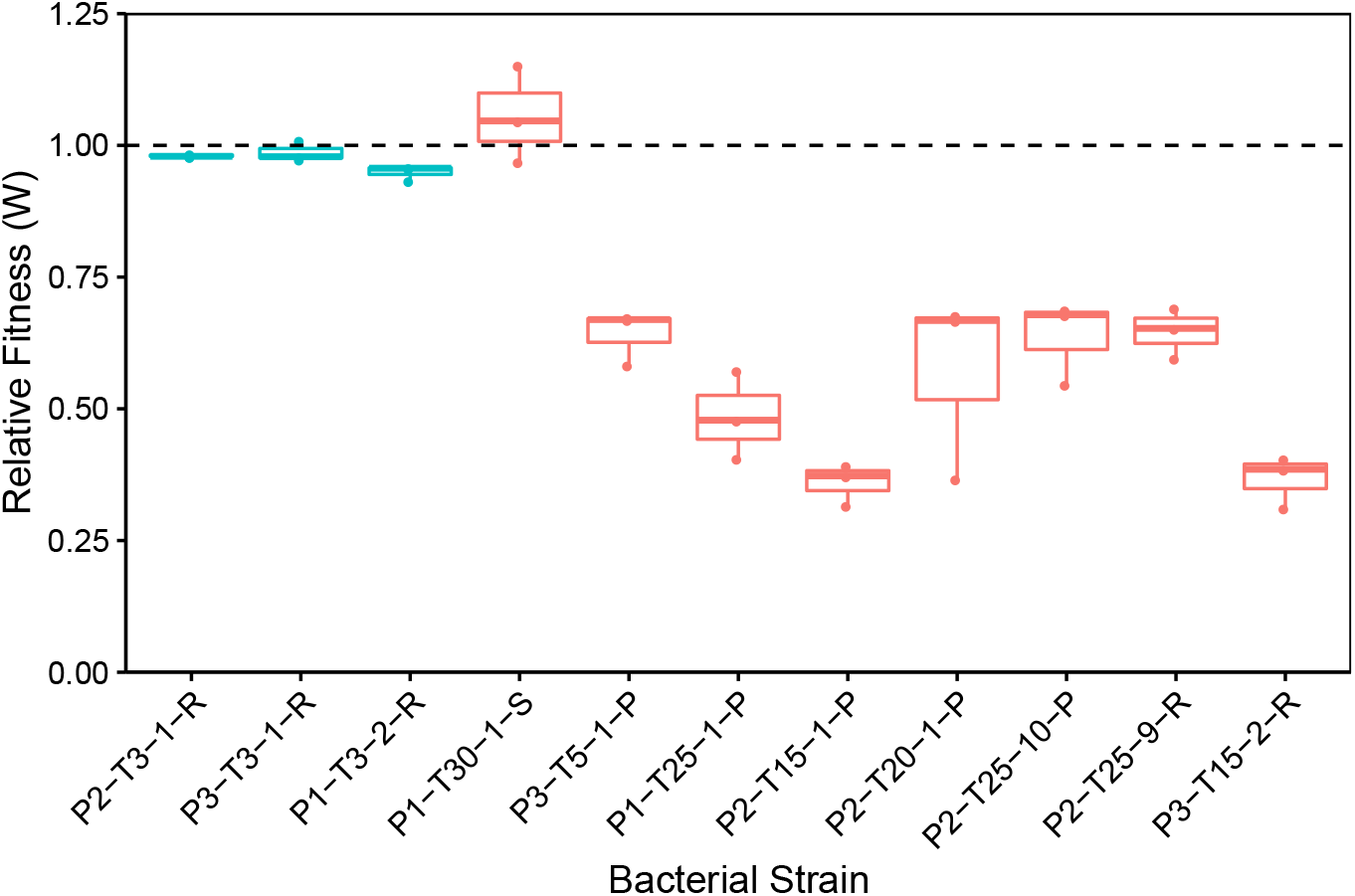
Fitness of coevolved bacterial isolates relative to their common ancestor, REL606. Isolates that coevolved with λunt are in teal and λtrn in red. Bacteria are labeled as in Fig. 3 except with treatment omitted. Mean relative fitness is significantly different between treatments with λunt 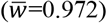 and λtrn (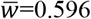, independent two-sample t-test, t=-8.19, p<<0.001). However, the relative fitness of isolates with partial and complete resistance to λtrn does not differ (p = 0.67). Strains are ordered firstly by treatment (λunt, λtrn) and then by increasing resistance according to PC1 (see Fig. 3).

### Phage Adaptation and Suppression of Resistance

Thus far, we have focused on the bacterial response to λunt and λtrn treatments. However, the ability of phages to evolve counter resistance to their hosts could have also contributed to differences in bacterial suppression and slowed the progression of resistance. To investigate whether phages evolved counter-defenses, we isolated a single phage from each population on the day we first detected that co-occurring bacteria had evolved resistance. We refer to co-occurring phage isolates as λunt+ and λtrn+ to differentiate them from their ancestors. We then conducted plate reader experiments to determine whether co-occurring phages (λunt+ or λtrn+) had evolved enhanced ability to suppress the earliest resistant bacteria to emerge from each population compared to ancestral phages (λunt or λtrn). We found that λunt+ phages were no better than λunt at suppressing resistant hosts (Fig. S5).

Contrastingly, all λtrn+ phages were better able to suppress co-occurring, resistant hosts than their ancestor, λtrn (Fig. 5). For Population 1, we compared the suppression of host P1-T25-1-P by λtrn and λtrn-P1-T25. We found greater suppression by λtrn-P1-T25, apparent visually (Fig. 5A), as well as quantitatively by comparing PC1 between replicate cultures (Fig. 5B, p=0.004, two-sample t-test). The same was true for λtrn+ phages from Population 2 (p=0.017, two-sample t-test, Fig. 5D, 5E) and Population 3 (p=0.034, two-sample t-test, Fig. 5G, 5H). For each population, comparisons were drawn from cultures where phage inoculum densities were not significantly different (*SI Appendix:* Phage Adaptation and Suppression of Resistance).

**Figure 5.**
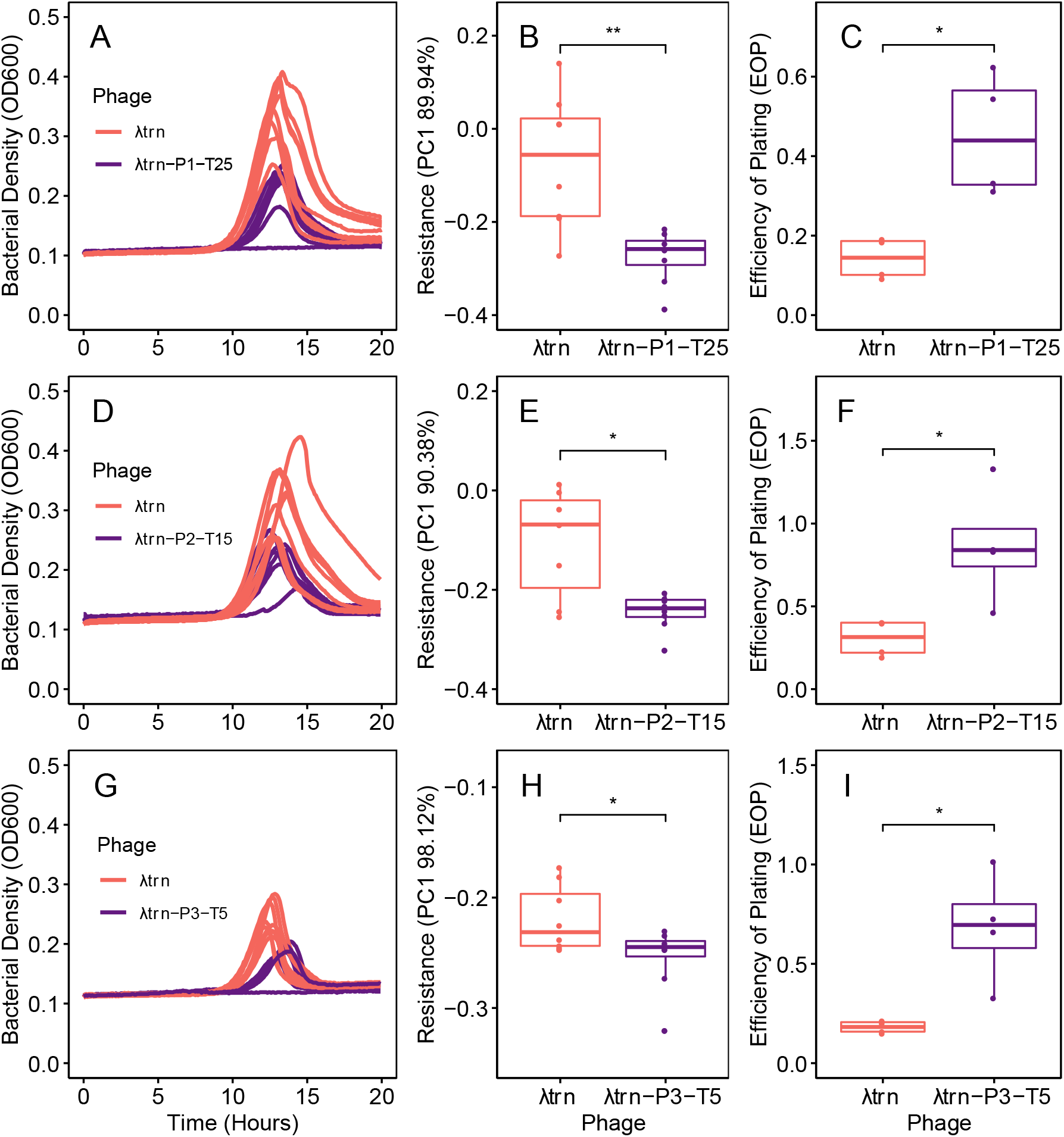
Coevolved λtrn+ isolates show adaptation and improved suppression of phage resistant bacteria. In all panels, comparisons are between ancestral phage λtrn (red) and a contemporary λtrn+ phage (purple) isolated from the same population and time-point as the resistant bacterial isolate. Panels A-C pertain to host P1-T25-1-P. Panels D-F pertain to host P2-T15-1-P. Panels G-I pertain to host P3-T5-1-P. Bacterial isolates are labeled as in Fig. 4. All hosts represent the first day at which λtrn resistance was detected. Panels A, D, G: Growth trajectories of replicate wells (n=4) inoculated with ∼1000 cells of the resistant host and either λtrn or λtrn+ phage. Panels B, E, H: Resistance of bacterial isolates to λtrn and respective λtrn+ phage (indicated by PC1, see Fig. S4). In all populations, bacterial isolates are less resistant to λtrn+ phages than to λtrn (Two-sample t-tests; B, p=0.004; E, p=0.017; H, p=0.034). Panels C, F, I: Efficiency of plating (EOP) of λtrn and λtrn+ phages on respective, resistant hosts compared to on REL606. In all populations, EOP is higher for λtrn+ phages than λtrn (C, p=0.022; F, p=0.045; I, p=0.037).

Support for these results also comes from efficiency of plating (EOP) assays. EOP, the ratio of plaques that a phage lysate forms on different bacteria, is often used to indicate how well phage are adapted to different hosts. All λtrn+ phages had higher EOP on resistant hosts than λtrn (Fig. 5C, 5F, 5I, p=0.022, p=0.045, p=0.037, respectively, two-sample t-tests). Although EOP alone is not indicative of improved suppression, these results support our conclusion that λtrn+ phages adapted to better suppress resistant hosts, whereas adaptation of λunt+ phages did not improve suppression. Two possibilities explain these results; λtrn may adapt to hosts more quickly and/or the kinds of resistance that evolved against λtrn may have been easier for the phage to counter. We did not investigate this further and do not know which contributed to our results.

### λtrn Adaptation via Recombination

At this point in the study, we explored how phage training enabled λtrn to gain its suppressive abilities (Table S1). Previously, it was reported that five mutations in λ’s (λunt’s) host recognition gene *J* gave it the ability to use the new receptor, OmpF (26). In the same gene, there is an enigmatic ∼1,300 bp region that experienced 49 mutations during phage training. These mutations appear to be the result of recombination between λ’s *J* gene (1545-2796) and REL606’s *ECB_00512* gene (882-2217). *ECB_00512* shares homology with *J* and *ECB_00512* is likely a genetic relict from past phage infection. The appearance of recombination in a gene under selection led us to question whether the recombination was adaptive. Using a previously reported phylogeny of the experiment in which the recombination evolved (35), we were able to deduce when the recombination occurred and engineer *J* alleles before and after the recombination. Next, we inserted the alleles and a genetic marker into λunt so that we could run competition experiments between pre- and post-recombination phages. We found that the recombination increased λunt’s ability to compete for wildtype (REL606), *malT-*, and *malT-OmpF-* host cells (Fig. 6; paired t-test of λ growth rates, p-values for wildtype, *malT-*, and *malT-OmpF-* were 0.002, 0.006, and 0.004, respectively). The recombination had the largest effect in competitions for *malT-ompF-* cells, which were the most difficult to infect and in which phage fitness doubled (ANOVA, p=0.005; Tukey’s Test p<0.05).

**Figure 6.**
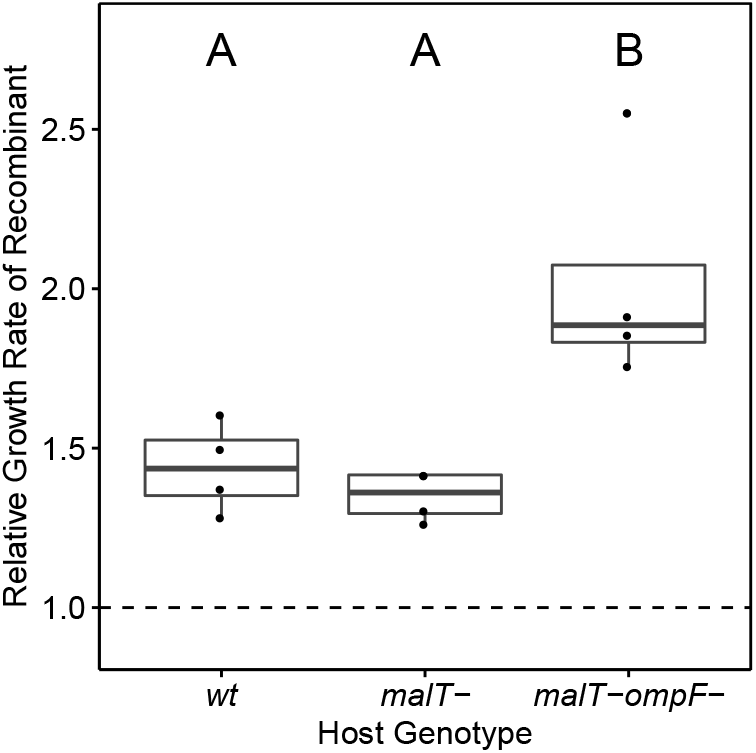
Increase in exponential growth rate of recombinant λ, relative to pre-recombination λ, when competing for three different *E. coli* host genotypes. Letters indicate groups that are statistically similar. The dashed line indicates where growth rate of the recombinant is equal to pre-recombination λ.

We find it remarkable that, through recombination, the relict DNA of a past infection provided information to improve infectivity of our contemporary phage. We expected that phage training worked entirely through the steady processes of coevolution, in which the phage makes incremental, reciprocal advances on its evolving host. However, recombination may accelerate training by transferring information from past phages that were well adapted to their host to new, naïve therapeutic phages. This may be an important mechanism for phage training success because therapeutic phages are typically isolated independently from the target bacterium (9,11,13) and may not be well adapted to these hosts. Many, if not most bacteria possess genes that originated from phage genomes (22,40) and phage recombination is frequently found in nature (41-43). Moreover, a previous study found that recombination with host encoded phages resulted in a host-range expansion (41), suggesting that recombination could be a dependable mechanism that enhances the effectiveness of phage training beyond the coevolutionary arms race. While the use of phage with the potential to recombine has generally been avoided due to the risk of transducing bacterial virulence or antibiotic resistance genes (43), here we show a potential benefit of phage-host recombination in phage training. If recombination is a concern during phage therapy, the recombination machinery could be left intact during training and then knocked out before treatment.

## Discussion

In phage therapy, the evolution of phage resistance is widespread and often renders phage therapeutics ineffective. Here, we evaluated the prospect of coevolutionary phage training as an approach to improve therapeutic efficacy by comparing untrained (λunt) and trained (λtrn) phages cocultured with a naïve target bacterium. We found that treatment with λtrn led to improved and prolonged suppression of host populations and that the evolution of λtrn resistance was delayed as compared to treatment with λunt. This prolonged suppression and delayed resistance was caused by several factors that all support coevolutionary phage training as an approach that could improve outcomes in phage therapy.

We found that it was more difficult for bacteria to evolve resistance to trained phage. Firstly, whereas individual mutations conferred complete resistance to λunt, mutations for λtrn resistance were ∼100× less common and only conferred partial resistance. Secondly, complete λtrn resistance required multiple mutations and both completely resistant bacteria isolated from the coevolution experiment had five putative resistance mutations. We believe the reduced ability of bacteria to evolve resistance to λtrn than to λunt was the greatest contributor to the stronger suppression and delayed resistance observed in the coevolution experiment. In part, these differences likely originate from the fact that λtrn can use two receptors (LamB and OmpF), whereas λunt can only use LamB. Phage resistance often evolves through augmentations of the receptors, and so deploying a phage with a backup receptor would be similar to using a drug cocktail where multiple mutations are required for bacteria to survive the treatment. It is unclear whether evolving to use a new receptor is a typical outcome of phage training; nonetheless, the use of dual receptor phages should be considered in developing future strategies for phage discovery (44), engineering (45,46), and directed evolution.

Thirdly, we found that λtrn resistance mutations carried large growth costs, regardless of whether they conferred partial or complete resistance, whereas mutations for λunt resistance carried little cost. In conjunction with the fact that, initially, resistance to λtrn is only partial, these large growth costs appear to have slowed the rate at which bacterial populations reached carrying capacity. Together, these factors may improve the ability of trained phage therapeutics, in combination with the host immune system, to resolve infections (47).

And fourthly, we found that previously trained phages were better able to counter the evolution of resistance during the experiment. In all populations, coevolved λtrn+ phages, isolated from the same time-point at which resistance was detected, were better able to suppress and had higher EOP on partially resistant hosts than their ancestor, λtrn. This was not true for coevolved λunt+ phages, which lost the ability to suppress resistant hosts. Although the role of phage coevolution in the efficacy of *in vivo* therapy has yet to be demonstrated, the ability of trained phages to coevolve with and counter host defenses may improve treatment outcomes.

Despite the close relatedness of the phages that we used, the effects of λtrn on the ecological and evolutionary dynamics of the microbial communities was vastly different from λunt. Flasks with λunt shifted from top-down (predator-controlled) to bottom-up (resource limited) systems where bacterial populations were limited by the availability of glucose in less than one day. With λtrn, flasks persisted as top-down systems in which phages suppressed bacterial populations below carrying capacity for several weeks. Moreover, the genetics underlying *E. coli*’s resistance evolution was very different. With untrained phage, bacteria acquired single, large-effect mutations that conferred complete resistance to λunt. However, with a trained phage, bacteria were constrained to evolve resistance through multiple mutations that each only yields partial resistance. These observations are fascinating from a basic biology perspective because they show how slight alterations in genomes and the evolutionary history of parasites can change their host’s ecology and evolution. From an applied standpoint, these observations provide proof of principle that directed evolution can vastly improve the effectiveness of phage therapeutics. Phage malleability is often lauded as a potential strength of phage therapy, and this is strongly supported by our findings.

Altogether, these results contribute to a small but growing body of work (23,48,49) in support of coevolutionary phage training as a means to improve the efficacy of phage therapeutics. Furthermore, they reveal the mechanisms by which this improved efficacy can be achieved: bacteria appear less able to resist trained phage and trained phage are more able to adapt to counter their hosts. We encourage efforts to investigate phage training on clinical strains of bacteria and to study the efficacy of trained phages in the treatment of animal model infections.

## Methods

### Strains

To study the effects of phage training on bacteria-phage coevolutionary dynamics we used *Escherichia coli* B strain REL606, as well as untrained (λunt) and trained (λtrn) strains of phage λ. For λunt, we used phage λcI26, a strictly lytic form of λ that has been characterized and compared with the λ reference genome by Meyer et al. (26). For λtrn, we used phage λ28-11, a descendant of λcI26 that was isolated from a plaque after coevolving with REL606 for 28 days (26,35). Notably, λtrn can infect through the native λ receptor, LamB, as well as a novel receptor, OmpF. We consider λtrn trained because it has previously coevolved with host REL606. Table S1 shows genetic comparisons between λtrn and λunt.

### Bacterial Suppression experiment

To determine whether trained (λtrn) phages would better suppress REL606 than untrained (λunt) phages, we inoculated six replicate 50 mL flasks with 10 mL modified M9 glucose media (M9-G, recipe in (26)) and ∼10^4^ cells. Of these flasks, 3 were inoculated with ∼10^8^ particles of λunt and 3 with ∼10^7^ particles of λtrn. Flasks were incubated at 37°C, shaking at 120 rpm. After 24 h, 100 μL of each community was transferred into new flasks with 10 mL of fresh media. Flasks were propagated for 30 days. Each day, aliquots were removed to estimate bacterial and phage densities, as well as to preserve communities for later analyses. For bacteria, aliquots were diluted in M9-G with 0.01M sodium citrate and plated on Luria-Bertani (LB) agar. For phages, 1 ml aliquots were treated with chloroform (5% v/v) and centrifuged (1 min at 15,000 x *g*) to extract lysates. Then, lysates were diluted in M9-G and 2 μL aliquots were spotted on infused soft agar (LB agar except with 0.8% w/w agar and inoculated with ∼10^8^ cells REL606 (26)). Lastly, starting on day 3, aliquots were preserved by freezing at -80°C in 15% v/v glycerol.

### Bacterial and Phage Isolation

To isolate bacteria, scrapes (∼2 μL) of preserved, frozen communities were streaked on LB agar plates and incubated overnight at 37°C. Then, colonies were isolated and streaked twice more to obtain clonal strains cleaned of phage. Strains were grown overnight at 37°C in M9-G and preserved by freezing. To isolate phages, scrapes of ice were suspended in tubes with 1 mL of M9-G. Then 10 μL was inoculated in molten (∼55°C) infused soft agar, poured over LB agar plates, and incubated overnight at 37°C. From these plates, plaques were isolated and plated in infused soft agar two more times. Then, plaques were picked into tubes with 4 mL of M9-G and ∼10^8^ cells REL606 and incubated overnight at 37°C. Lastly, lysates were extracted and preserved as described above.

### Emergence of Resistance

To determine when resistance evolved to λunt and λtrn, we used 12 bacterial isolates from each community across various days of the coevolution experiment. We conducted resistance assays in liquid using 96-well plates and a plate reader (Tecan Sunrise). For each isolate, overnight cultures in LB were diluted and ∼10^3^ cells inoculated into wells containing either 200 μL of LBM9 or 150 μL of LBM9 with 50 μL of 0.22 μm filtered lysate prepared in LBM9 (∼10^8^ phage particles, recipe in (27)). Optical density (OD) at wavelength 600 nm was read from plates over 20 h of incubation, shaking at 37°C. We used LBM9 instead of M9-G for plate reader assays because experiments in M9-G were unable to distinguish between sensitivity and resistance against λunt. In M9-G, the native λ receptor, LamB, is downregulated and λunt does not adsorb to nor kill cells efficiently (50,51). In contrast, LBM9 allowed us to distinguish sensitive, partial, and complete resistance.

### Estimating Resistance Mutation Rate

To estimate REL606’s mutation rate for resistance (μ_r_) against λunt and λtrn, we conducted Luria-Delbruck fluctuation tests in liquid culture. REL606 was grown in LB overnight, diluted, inoculated into independent isogenic cultures (∼10^3^ cells per well) in a 96-well plate, and grown again overnight. Cultures were diluted and 10 μL inoculated into a new plate with 150 μL of LBM9 and 50 μL of filtered LMB9 lysate (∼10^8^ phage particles) in each well. To estimate μ_r_ to λunt, fluctuation tests were conducted with 10^4^- and 10^5^-cell inoculums. To estimate μ_r_ to λtrn, we used 10^7^-cell inoculums. Plates were read for 20 h while shaking at 37°C. Wells with no observable growth were deemed sensitive. The P_0_ method (P_0_ = *e*^μr*(N-No)^) was used to calculate μ_r_, where P_0_ is the proportion of wells that were sensitive, N is the number of cells inoculated in each well, and No is the number of cells used to start cultures (∼10^3^ cells). To screen for false positives, bacteria were isolated from wells showing resistance and tested in 96-well plates as described above.

### Bacterial and phage genomics

To extract genomes, bacteria were streaked from frozen stocks onto LB agar and incubated overnight. Colonies were picked into respective growth media (M9-G for coevolution, LB for fluctuation test), incubated overnight, and genomes were extracted using an Invitrogen PureLink Genomic DNA Mini Kit. For phages, lysates were prepared by incubating ∼2 μL from frozen stocks with ∼10^8^ cells of REL606 overnight in M9-G and then extracted using 0.22 μm filters. To extract genomes, 500 μL of lysate was transferred to 1.5 mL microcentrifuge tubes with 50 μL of DNase buffer (10×), 10 U of DNase I, and 1 μL of 100 mg/mL RNaseA and incubated for 30 min at room temperature. Then, 20 μL of 0.5 M EDTA, 2.5 U of proteinase K, and 25 μL of 10% SDS were added and tubes incubated at 55°C for 1 h. Lastly, phenol chloroform extractions were conducted to purify the phage DNA [52]. The resulting supernatant and an equal volume of 100% EtOH were added to spin columns from a Purelink Genomic DNA Mini Kit and then washed and eluted according to the manufacturer’s instructions. Finally, extracted bacterial and phage genomes were sent to the Microbial Genome Sequencing Center (MiGS) where they were indexed and sequenced on an Illumina NextSeq 550. Sequences were analyzed using *breseq* (version 0.35.0), a computational pipeline for analyzing short-read re-sequencing data [33].

### Genetics of Resistance

To investigate the relationship between the number of resistance mutations and the level of resistance in bacterial isolates, we first identified resistance mutations as those occurring in genes that are involved in λ infection or that evolved in multiple populations, fluctuation tests, or previous coevolution experiments. Then, we created a scale indicative of resistance by taking the difference in OD trajectories of bacterial cultures growing without and with phage and conducting principal component analysis (PCA) on these OD difference vectors in R (version 3.6.1, (53)). PCA revealed that PC1 explained 92.25% of the variance and discriminated between sensitive, partial, and complete resistance along a continuous scale. Using PC1 values as resistance scores, we used R to implement linear models (PC1∼number of resistance mutations) testing the relationship between mutations and resistance against each phage.

### Costs of Phage Resistance

Competitions were conducted between coevolved bacterial isolates and an *ara*+ marked REL606 ancestor. To obtain the marked ancestor, ∼10^8^ cells of REL606 (*ara-*) were plated on minimal-arabinose agar plates (5.34 g potassium phosphate dibasic anhydrous, 2 g potassium phosphate monobasic anhydrous, 1 g ammonium sulfate, 0.57 g sodium citrate dihydrate, 16 g agar, 4 g arabinose per liter of water and supplemented to a final concentration of 1 mM magnesium sulfate, 0.0002% w/v thiamine, and 0.0002% w/v biotin). After 2 days of incubation at 37°C, colonies were picked and streaked twice on tetrazolium-arabinose indicator agar (Tet-ara) plates (10 g tryptone, 1 g yeast extract, 5 g sodium chloride, 16 g agar, 10 g arabinose per liter of water and supplemented to a final concentration of 0.005% tetrazolium indicator dye TTC) to confirm marker presence and obtain isogenic stocks. To initiate competitions, colonies on Tet-ara plates were picked and incubated for 24 h as in the coevolution experiment. The next day, coevolved strains and the marked competitor were inoculated 1:9 or 1:99 to a final volume of 100 μL in fresh media. Upon inoculation, flasks were mixed, and an aliquot was diluted and plated on Tet-ara plates to enumerate initial densities (T_0_) of each strain. After competing at 37°C for 24 h, aliquots were again diluted and plated on Tet-ara plates for final densities (T_F_). Finally, relative fitness (W) was calculated for each strain where W = M_A_/M_B_ and where M_A_ = ln [T_F_/T_0_] of the coevolved strain and M_B_ = ln [T_F_/T_0_] of the marked ancestor.

### Phage Adaptation and Suppression of Resistance

First, coevolved phages were isolated from frozen communities as described above. Then, plate reader experiments were conducted as described in Emergence of Resistance except with the following changes: phage lysates were prepared on REL606 but 96-well plates were inoculated with representative resistant bacteria. Ancestral (λunt, λtrn) and coevolved phages (λunt+, λtrn+) were inoculated at various dilutions and were also enumerated by spotting 2 μL of the lysate dilution series on soft agar plates infused with REL606 in quadruplicate. Then, PCA was conducted on OD growth trajectories of bacteria-phage cocultures where phage inoculum densities were not significantly different between treatments. We also spotted 2 μL of the dilution series on resistant isolates to calculate efficiency of plating (EOP) on resistant hosts where EOP = [titer on resistant host] / [titer on REL606].

### Phage Adaptation via Recombination

We constructed 4 phage strains from λunt with pre- and post-recombination *J* alleles, both with and without a *lacZ* genetic marker, in order to isolate the effect of the *J* mutations. Using a previously constructed phylogeny of the experiment in which the recombination evolved (35), we determined that the recombination occurred within a *J* allele that had already evolved 7 mutations (C2999T, A3034G, T3230, C3310, G3319A, T3321A, A3364T). Therefore, we constructed alleles with these 7 mutations with and without the recombination. Fortunately, the 7 mutations do not overlap with the recombination, so we were able to PCR amplify DNA fragments with either the 7 mutations or the entire gene using the primers in Table S2. These fragments were ligated into a plasmid using an Invitrogen TA Cloning™ Kit and the plasmids were used to transform *E. coli* DH5α. λunt was modified by infecting these cells and then screening for particles that could use the non-native OmpF receptor as a result of incorporating the *J* alleles from the plasmid. We constructed two independent versions of each phage genotype and the incorporation of the desired alleles was sequence verified. Next, we added a *lacZ* genetic marker which causes the phage to produce blue plaques (54,55). Head-to-head competitions were performed in conditions identical to the original evolution experiment (26). Competitions for 3 different hosts were evaluated; wildtype (REL606), *malT-* (REL606 with a 25 bp duplication in *malT*), and *malT-ompF-* (*malT-* with a single base substitution resulting in an amino acid change (N52K) in *ompF* (full genome in (26))). Four replicates were performed for each competition. Half of the competitions were conducted with pre-recombination genotypes marked with *lacZ* and the other half with the post-recombination genotype marked. Each flask was initiated with ∼10^6^ particles of each λ genotype and ∼10^8^ cells. Initial and final phage densities were enumerated by counting plates on X-gal agar indicator plates (recipe in (54)).

## Acknowledgements

We thank members Sarah Medina and Elijah Horwitz of the J.R.M. laboratory for productive discussions and feedback on this study. This work was supported by the United States – Israel Binational Science Foundation: 2017056 and the National Institutes of Health (NIH) Grant R01: GM088344.

## Notes

### Competing Interest Statement

The authors have declared no competing interest.

