## Supplementary material for "Coevolutionary phage training leads to greater bacterial suppression and delays the evolution of phage resistance": SI Appendix

**Table S1.** Genomic differences between untrained phage λunt (λcI26) and trained phage λtrn. For genomic differences between λunt and the λ reference genome (GenBank: NC_001416) see Meyer et al. 2012 (1). Synonymous mutations are in green and nonsynonymous mutations in blue. Mutations from position 17,049 to 17,868 evolved via recombination with a relict prophage in the genome of REL606 (1).

| Genome position | Mutation | Type | Amino acid change | Gene | Product |
|---|---|---|---|---|---|
| 11,451 | C → T | Substitution | A304V | *H* | Tail component |
| 17,049 | C → T | Recombination | S515S | *J* | Tail: host specificity protein |
| 17,055 | T → C | Recombination | G517G | *J* | Tail: host specificity protein |
| 17,059 | G → A | Recombination | A519T | *J* | Tail: host specificity protein |
| 17,081 | +G | Recombination | — | *J* | Tail: host specificity protein |
| 17,082 | A → C | Recombination | G526G | *J* | Tail: host specificity protein |
| 17,085 | Δ1 bp | Recombination | — | *J* | Tail: host specificity protein |
| 17,088 | C → G | Recombination | G528G | *J* | Tail: host specificity protein |
| 17,090 | A → G | Recombination | N529S | *J* | Tail: host specificity protein |
| 17,136 | A → G | Recombination | V544V | *J* | Tail: host specificity protein |
| 17,160 | T → C | Recombination | G552G | *J* | Tail: host specificity protein |
| 17,183 | A → G | Recombination | E560G | *J* | Tail: host specificity protein |
| 17,200 | C → T | Recombination | L566L | *J* | Tail: host specificity protein |
| 17,211 | A → C | Recombination | R569R | *J* | Tail: host specificity protein |
| 17,280 | G → A | Recombination | V592V | *J* | Tail: host specificity protein |
| 17,328 | A → C | Recombination | E608D | *J* | Tail: host specificity protein |
| 17,334 | T → C | Recombination | S610S | *J* | Tail: host specificity protein |
| 17,343 | G → A | Recombination | V613V | *J* | Tail: host specificity protein |
| 17,391 | T → C | Recombination | T629T | *J* | Tail: host specificity protein |
| 17,409 | T → C | Recombination | Y635Y | *J* | Tail: host specificity protein |
| 17,421 | G → C | Recombination | A639A | *J* | Tail: host specificity protein |
| 17,424 | A → C | Recombination | R640R | *J* | Tail: host specificity protein |
| 17,430 | C → T | Recombination | D642D | *J* | Tail: host specificity protein |
| 17,433 | A → G | Recombination | T643T | *J* | Tail: host specificity protein |
| 17,457 | T → C | Recombination | S651S | *J* | Tail: host specificity protein |
| 17,466 | C → T | Recombination | L654L | *J* | Tail: host specificity protein |
| 17,469 | T → C | Recombination | R655R | *J* | Tail: host specificity protein |
| 17,478 | +GG | Recombination | — | *J* | Tail: host specificity protein |
| 17,487 | C → T | Recombination | D661D | *J* | Tail: host specificity protein |
| 17,494 | A → C | Recombination | S664R | *J* | Tail: host specificity protein |
| 17,502 | G → A | Recombination | R666R | *J* | Tail: host specificity protein |
| 17,535 | A → T | Recombination | T677T | *J* | Tail: host specificity protein |
| 17,547 | G → A | Recombination | T681T | *J* | Tail: host specificity protein |
| 17,556 | G → T | Recombination | A684A | *J* | Tail: host specificity protein |
| 17,586 | G → A | Recombination | A694A | *J* | Tail: host specificity protein |
| 17,613 | T → G | Recombination | D703E | *J* | Tail: host specificity protein |
| 17,652 | A → G | Recombination | A716A | *J* | Tail: host specificity protein |
| 17,659 | 2bp → CA | Recombination | — | *J* | Tail: host specificity protein |
| 17,673 | G → A | Recombination | T723T | *J* | Tail: host specificity protein |
| 17,679 | C → G | Recombination | G725G | *J* | Tail: host specificity protein |
| 17,721 | C → T | Recombination | D739D | *J* | Tail: host specificity protein |
| 17,759 | A → G | Recombination | Q752R | *J* | Tail: host specificity protein |
| 17,775 | A → G | Recombination | R757R | *J* | Tail: host specificity protein |
| 17,788 | +CA | Recombination | — | *J* | Tail: host specificity protein |
| 17,795 | Δ2 bp | Recombination | — | *J* | Tail: host specificity protein |
| 17,805 | T → C | Recombination | G767G | *J* | Tail: host specificity protein |
| 17,862 | C → T | Recombination | Y786Y | *J* | Tail: host specificity protein |
| 17,868 | T → C | Recombination | Y788Y | *J* | Tail: host specificity protein |

| | | | | | |
|---|---|---|---|---|---|
| 18,285 | C → A | Recombination | D927E | *J* | Tail: host specificity protein |
| 18,297 | 4bp → ATAT | Recombination | — | *J* | Tail: host specificity protein |
| 18,503 | C → T | Substitution | A1000V | *J* | Tail: host specificity protein |
| 18,538 | A → G | Substitution | S1012G | *J* | Tail: host specificity protein |
| 18,814 | C → T | Substitution | H1104Y | *J* | Tail: host specificity protein |
| 18,823 | G → A | Substitution | D1107K | *J* | Tail: host specificity protein |
| 18,825 | T → A | Substitution | D1107K | *J* | Tail: host specificity protein |
| 18,868 | A → T | Substitution | I1122F | *J* | Tail: host specificity protein |
| 19,260 | T → C | Substitution | L99P | *lom* | Outer host membrane |
| 20,661 | A → G | Substitution | K338E | *Orf-401* | Tail fiber protein |
| 45,176 | $(G)_{5\rightarrow 6}$ | Insertion | — | *Orf-64* | Hypothetical prot./anti-holin |

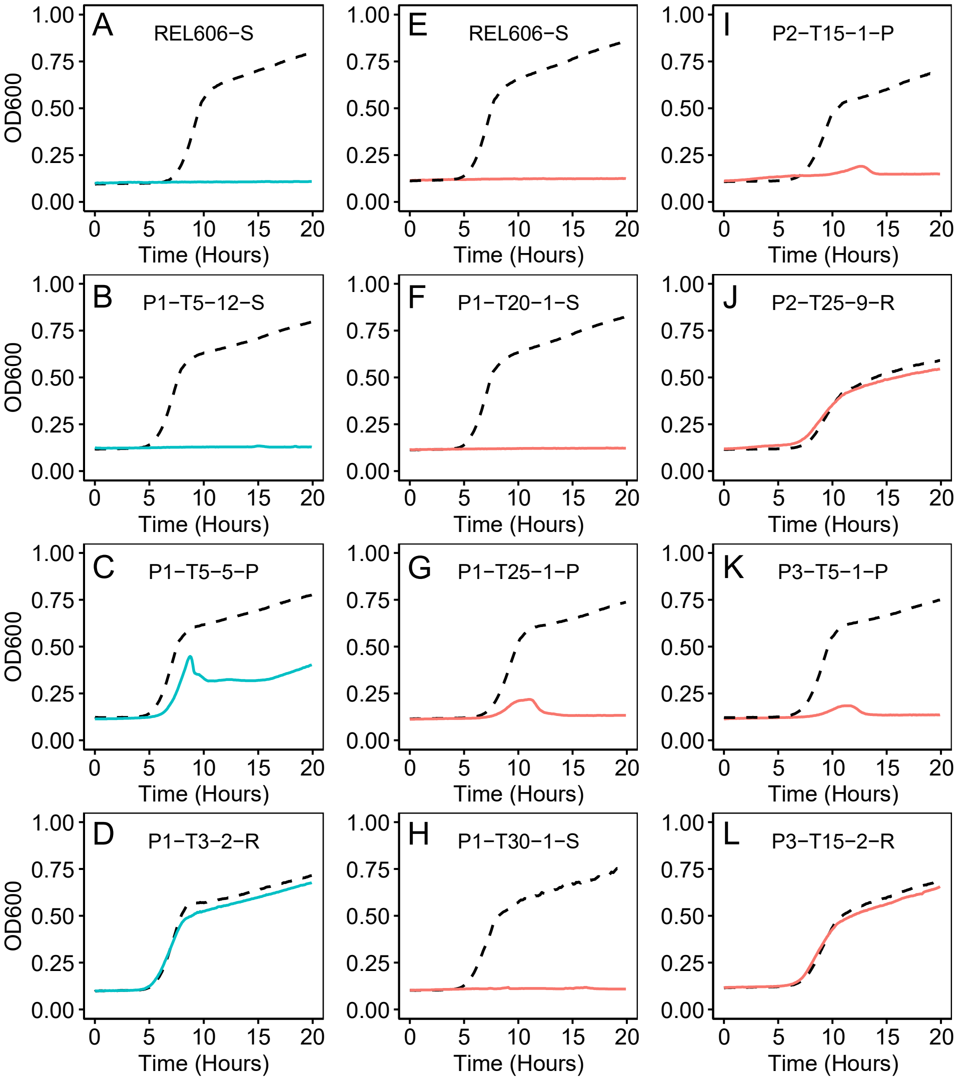


**Figure S1.** Diversity of resistance profiles determined from OD growth trajectories of bacterial isolates growing without (black, dotted) and with phage (solid, teal for λunt, red for λtrn) for 20 h. Plot titles indicate the bacterial isolate and resistance status. Apart from the ancestor REL606, isolates are labeled as [Population]-[Day Isolated]-[Isolate #]-[Resistance Status], where S = Sensitive, P = Partial Resistance, and R = Complete Resistance. Isolates were deemed sensitive if no growth was observed in the presence of phage, partially resistant if there was growth and inhibition, and completely resistant if there appeared to be no inhibition of growth by phage. Panels A-D correspond with phage λunt and panels E-L with λtrn.

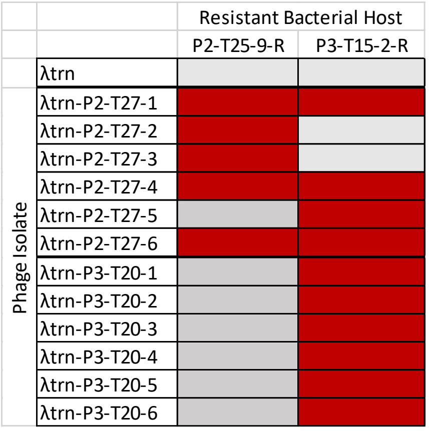


**Figure S2.** Infectivity of λtrn and descendant phages on bacterial hosts isolated from the coevolution experiment. Hosts (columns, labeled as in Fig. S1) were completely resistant to λtrn due to *malT* and *OmpF* mutations (see Fig. S3). Lysates of λtrn and descendant phage isolates were spotted onto host infused LB agar. Fill indicates resistance/infectivity (grey = resistant, red = sensitive).

**λtrn Resistance Mutations.**

The mechanisms by which some mutations affected λtrn resistance were unclear. All bacterial isolates from the coevolution experiment possessed a 12,090 bp deletion comprising 14 unannotated genes. Moreover, all populations (but not all isolates) were also found to have a 4 bp CCAG duplication in the coding sequence of another unannotated gene. And thirdly, the λtrn-sensitive isolate P1-T30-1 had a 777 bp IS element excision of loci *insB-22*, *insA-22* and *ECB_02825* that have previously been found to confer λtrn resistance through epistasis with mutations in *malT* (2). However, no *malT* mutations were found in any of the 12 strains isolated from day 30. These results beg the question: How did these bacteria reach high densities in flasks with λtrn if they were sensitive? The mechanism by which the 777 bp deletion interacts with mutations in *malT* is unknown and it is possible that this deletion also interacts with mutations at other loci. Although we were unable to detect growth of day-30 isolates when cocultured with λtrn, it is likely that these isolates are more resistant than their ancestor REL606—but that our assay was not sensitive enough to discriminate these differences.

Lastly, we point out two instances where λtrn recombined into the host genome. In λtrn Population 2, the completely resistant isolate P2-T25-9 had 18 mutations in *ybcD*, a predicted prophage replication protein fragment gene, from positions 539,643 to 539,915. Additionally, the completely resistant isolate P3-T15-2 from Population 3 had 4 mutations in *ybcT*, a phage lysis gene, from positions 549,746 to 549,772. BLASTX of mutant sequences to REL606 and λ references genomes revealed a 100% match with λ, whereas they showed 89% and 96.88% identity, with REL606, respectively, confirming that these sequences originated from the phage. Although both recombinations from phage were found in bacteria that evolved complete resistance, we have no evidence that these recombinations carry any selective benefit. More likely, recombination occurred due to the endogenous lambda red system which recombines homologous sequences at a high rate.

| Genome position | P1−T3−2−R | P2−T3−1−R | P3−T3−1−R | P1−T5−5−P | P1−T25−1−P | P1−T30−1−S | P2−T15−1−P | P2−T20−1−P | P2−T25−10−P | P2−T25−9−R | P3−T5−1−P | P3−T15−2−R | JB42−P | JB43−P | JB47−R | Mutation | Annotation | Gene |
|---|---|---|---|---|---|---|---|---|---|---|---|---|---|---|---|---|---|---|
| 12,965 | | | | | | | | | | | | | | | ■ | 15 bp duplication* | coding | *dnaK* |
| 37,281 | | | | | | | | ■ | | | | | | | | C→A | A798A (GCC→GCA) | *carB* |
| 112,876 | | | | | | | | | | | | ■ | | | | $(T)_{5\rightarrow 6}$ | coding | *secA* |
| 242,025 | | | | | | | ■ | ■ | ■ | ■ | | | | ■ | | Δ21,535 bp | between IS*1* | *ECB_00212–phoE* |
| 248,651 | | | | | | | | | | | | | ■ | | ■ | C→A | H180N | *lpcA* |
| 539,643 | | | | | | | | | | ■ | | | | | | G→T | pseudogene | *ybcD* |
| 539,649 | | | | | | | | | | ■ | | | | | | C→T | pseudogene | *ybcD* |
| 539,661 | | | | | | | | | | ■ | | | | | | G→C | pseudogene | *ybcD* |
| 539,664 | | | | | | | | | | ■ | | | | | | C→T | pseudogene | *ybcD* |
| 539,670 | | | | | | | | | | ■ | | | | | | T→C | pseudogene | *ybcD* |
| 539,673 | | | | | | | | | | ■ | | | | | | C→A | pseudogene | *ybcD* |
| 539,679 | | | | | | | | | | ■ | | | | | | A→G | pseudogene | *ybcD* |
| 539,681 | | | | | | | | | | ■ | | | | | | +T | pseudogene | *ybcD* |
| 539,686 | | | | | | | | | | ■ | | | | | | 2 bp→A | pseudogene | *ybcD* |
| 539,694 | | | | | | | | | | ■ | | | | | | C→T | pseudogene | *ybcD* |
| 539,698 | | | | | | | | | | ■ | | | | | | C→A | pseudogene | *ybcD* |
| 539,721 | | | | | | | | | | ■ | | | | | | A→C | pseudogene | *ybcD* |
| 539,827 | | | | | | | | | | ■ | | | | | | T→G | pseudogene | *ybcD* |
| 539,879 | | | | | | | | | | ■ | | | | | | G→T | intergenic | *ybcD* / *renD* |
| 539,891 | | | | | | | | | | ■ | | | | | | A→G | intergenic | *ybcD* / *renD* |
| 539,893 | | | | | | | | | | ■ | | | | | | A→C | intergenic | *ybcD* / *renD* |
| 539,907 | | | | | | | | | | ■ | | | | | | G→C | intergenic | *ybcD* / *renD* |
| 539,915 | | | | | | | | | | ■ | | | | | | T→C | intergenic | *ybcD* / *renD* |
| 549,746 | | | | | | | | | | | | ■ | | | | C→T | R57R (CGC→CGT) | *ybcT* |
| 549,764 | | | | | | | | | | | | ■ | | | | A→G | A63A (GCA→GCG) | *ybcT* |
| 549,767 | | | | | | | | | | | | ■ | | | | G→C | L64L (CTG→CTC) | *ybcT* |
| 549,772 | | | | | | | | | | | | ■ | | | | A→C | E66A (GAA→GCA) | *ybcT* |
| 787,879 | | | | | ■ | ■ | ■ | ■ | ■ | ■ | ■ | ■ | | | | Δ12,090 bp | | *ECB_00726–ECB_00739* |
| 1,003,024 | | | | | | | | ■ | ■ | ■ | | | | | | C→A | D351Y (GAC→TAC) | *ompF* |
| 1,003,277 | | | | | | | | | | | | ■ | | | | A→C | F266L (TTT→TTG) | *ompF* |
| 1,117,882 | | ■ | | | | | | | | | | | | | | G→A | intergenic | *csgD* / *csgB* |
| 1,230,063 | | | | | | | | ■ | | | | | | | | $(T)_{8\rightarrow 9}$ | pseudogene | *hlyE* |
| 1,893,290 | | | | | | | | ■ | ■ | ■ | | | | | | A→C | L35* (TTA→TGA) | *prc* |
| 2,103,918 | | | | | ■ | ■ | | ■ | ■ | ■ | | ■ | | ■ | | $(CCAG)_{7\rightarrow 8}$ | coding | *ECB_01992* |
| 2,183,354 | | | | | | | | | | | | | | ■ | | A→G | E91E (GAA→GAG) | *cdd* |
| 3,023,945 | | | | | | ■ | | | | | | | | | | Δ777 bp | | *insB-22–ECB_02825* |
| 3,481,793 | | | | ■ | | | | | | | | | | | | A→T | I37F (ATC→TTC) | *malT* |
| 3,482,495 | | | | | | | | | | ■ | | | | | | Δ20 bp | coding | *malT* |
| 3,482,706 | ■ | ■ | ■ | | | | | | | | | | | | | 25 bp duplication† | coding | *malT* |
| 3,734,067 | | | | | ■ | | | | | | | | | | | Δ1 bp | coding | *gumD* |
| 3,736,304 | | | | | | | | | | | ■ | ■ | | | | Δ4,894 bp | IS*1*-mediated | *waaC–waaT* |
| 4,141,100 | | | | | | | | | | | ■ | | | | | G→T | R58L (CGT→CTT) | *yijC* |

**Figure S3.** Mutations of representative bacterial isolates. Teal and red correspond with isolates from λunt and λtrn coevolution treatments, respectively. Orange corresponds with isolates from fluctuation tests. Coevolution isolates are labeled as in Fig. S1. Mutations were classified as putatively resistant if they occurred in genes 1) known to be involved in λ infection or 2) evolved in multiple populations, fluctuation tests, or previous coevolution experiments (yellow rows). *(AAGCGGCAGAAAAAG) †(AGTGGGAACTGGCGGCGGAGCTGCC).

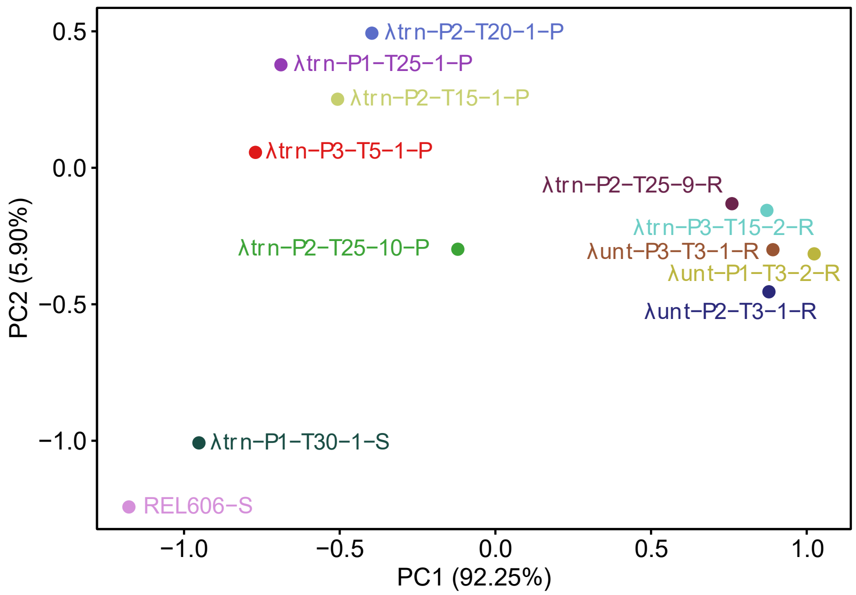


**Figure S4.** Principal Component Analysis (PCA) of the differences in OD trajectories of representative bacterial isolates growing without and with phages in plate reader experiments. PC1 distinguishes between sensitive, partial, and complete resistance against ancestral phages from respective coevolution treatments (λunt or λtrn). Apart from the ancestor REL606, isolates are labeled adjacent to points as in Fig. S1 and including their treatment.

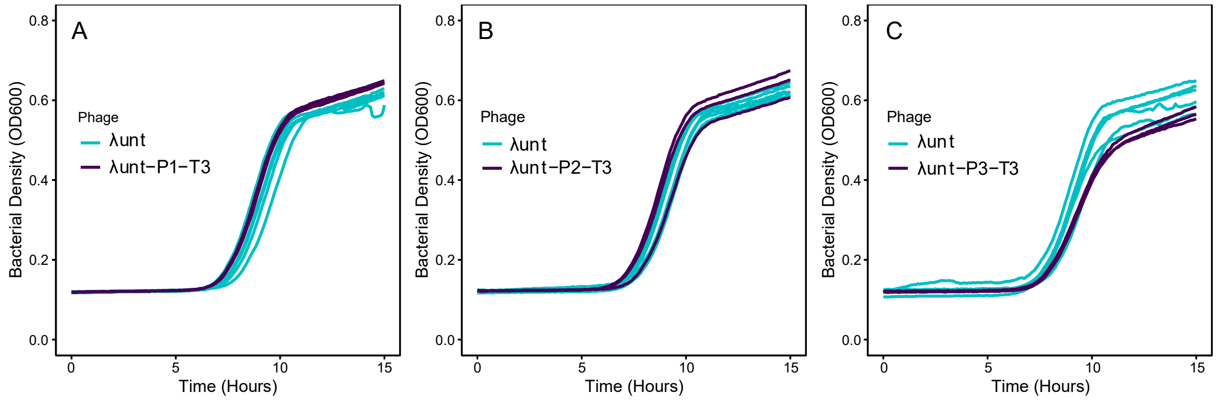


**Figure S5.** Coevolved λunt+ phages do not show improved suppression of resistant bacterial isolates. Growth trajectories of replicate wells were inoculated with ~1000 cells of the resistant host and either λunt (teal, n=6) or a contemporary λunt+ phage (purple, n=3) from the same population and time-point as the host. Panel A pertains to host P1-T3-2-R. Panel B pertains to host P2-T3-1-R. Panel C pertains to host P3-T3-1-R. All hosts represent the first day at which λunt resistance was detected.

**Phage Adaptation and Suppression of Resistance.** For λtrn Population 1, comparisons were drawn from cultures where phage inoculums were not significantly different (p=0.09, two-sample t-test) and the mean density of λtrn ($\bar{x}=1.17\times10^8$ pfu/mL) was higher than λtrn-P1-T25 ($\bar{x}=6.25\times10^7$ pfu/mL). For Population 2, λtrn-P2-T15 led to greater suppression (p=0.02, two sample t-test, Fig. 5D, 5E) when phage inoculums were not significantly different (p=0.22, two-sample t-test) and the mean density of λtrn-P2-T15 ($\bar{x}=8.13\times10^7$ pfu/mL) was higher than λtrn ($\bar{x}=5.39\times10^7$ pfu/mL). We note that when mean density of λtrn was higher than λtrn-P2-T15 (albeit not significantly, p=0.24) we found no difference in suppression. Lastly, for Population 3, λtrn+ phage led to greater suppression (p=0.034, two sample t-test, Fig. 5G, 5H) when the mean density of λtrn ($\bar{x}=3.19\times10^8$ pfu/mL) was higher than for λtrn-P3-T5 ($\bar{x}=9.69\times10^7$ pfu/mL). Notably, when significantly more λtrn was used ($\bar{x}=6.38\times10^8$ pfu/mL compared to $\bar{x}=3.88\times10^8$ pfu/mL, p=0.02, two-sample t-test), λtrn-P3-T5 still showed significantly more suppression of host P3-T5-1-P (p=0.02, two-sample t-test).

**Table S2.** Primers used to amplify segments of λ gene *J* for engineering alleles with the recombination.

| Primer ID | Primer Position | Sequence (5’ → 3’) |
|---|---|---|
| J Forward | 15 bp upstream | CTGCGGGCGGTTTTGTCATT |
| 2906 J Forward | Intergenic (2906) | CTCCGGGACGCTCAGTA |
| J Reverse | 318 bp downstream | ACGTATCCTCCCCGGTCATCACT |
